# Neuronal Hyperexcitability: A Key to Unravelling Hippocampal Synaptic Dysfunctions in Lafora Disease

**DOI:** 10.1101/2025.04.23.650182

**Authors:** Cinzia Costa, Laura Bellingacci, Jacopo Canonichesi, Valentina Imperatore, Anna Aurora Taddei, Luis Zafra-Puerta, Nerea Iglesias-Cabeza, Paolo Prontera, Andrea Mancini, Massimiliano Di Filippo, Alessandro Tozzi, Marta Barzasi, Fabrizio Gardoni, Marina P. Sánchez, José M. Serratosa, Lucilla Parnetti, Miriam Sciaccaluga

**Affiliations:** Section of Neurophysiopathology S.M. della Misericordia Hospital, Section of Neurology and Laboratory of Experimental Neurology, Department of Medicine and Surgery, University of Perugia, Perugia 06132, Italy; Section of Physiology and Biochemistry, Department of Medicine and Surgery, University of Perugia, Perugia 06132, Italy; Laboratory of Experimental Neurology, Department of Medicine and Surgery, University of Perugia, Perugia 06132, Italy; Laboratory of Neurology, Instituto de Investigación Sanitaria-Fundación Jiménez Díaz, Universidad Autónoma de Madrid (IIS-FJD, UAM), 28040 Madrid, Spain; PhD Program in Neuroscience, Universidad Autónoma de Madrid-Cajal Institute, Madrid, Spain 28029; Fondazione Malattie Rare Mauro Baschirotto BIRD Onlus, Longare (VI), Italy; Medical Genetics and Rare Diseases Unit, Maternal-infantile Department, S.M. della Misericordia Hospital, Perugia, Italy; Section of Neurology, Department of Medicine and Surgery, University of Perugia, Perugia, Italy; Department of Pharmacological and Biomolecular Sciences, University of Milano, Milan, Italy

**Keywords:** Lafora Disease, Neuronal Hyperexcitability, Neurodegeneration, Synaptic Plasticity, Hippocampal Synaptic Dysfunction, Cannabidiol, Therapeutic Window

## Abstract

Lafora disease (LD) is a rare progressive disorder caused by mutations in the EPM2A or EPM2B genes, characterized by the accumulation of Lafora bodies, drug-resistant epilepsy, and cognitive decline. To investigate the early molecular mechanisms of LD, we studied electrophysiological changes in the dentate gyrus (DG) of the Epm2aR240X knock-in mouse model at various ages. Electrophysiological recordings measured neuronal membrane properties, epileptic-like activity, epileptic thresholds, and synaptic plasticity in Epm2aR240X mice at 1, 3, and 12 months. We also employed PAS diastase staining, immunofluorescence, and Western blotting to detect Lafora bodies, amyloid beta deposition, and glutamate receptor subunit expression.

Epileptic-like activity began at 1 month and intensified with age. Aberrant long-term potentiation (LTP) appeared at 3 months and worsened by 12 months. Notably, cannabidiol (CBD) treatment reduced excitability and restored LTP in older mice, suggesting its potential therapeutic value. These findings indicate that network hyperexcitability is an early event in LD, highlighting a therapeutic window for interventions like CBD.

## Introduction

Lafora disease (LD, OMIM #254780) presents a challenge within the spectrum of rare, progressive neurodegenerative diseases, striking without gender bias (1). It originates from mutations in the *EPM2A* and *EPM2B* genes (2–6), which encode laforin and malin, two enzymes crucial for regulating glycogen metabolism. These mutations lead to the hallmark features of LD, including medication-resistant epilepsy and neurodegeneration (7). LD is marked by the buildup of Lafora bodies (LBs), insoluble polyglucosan inclusions found in both cerebral and peripheral tissues (8–10). The disease typically manifests during adolescence in previously healthy individuals. Beginning with generalised tonic-clonic seizures, it progresses to cognitive decline, dementia, and increasing seizure frequency, ultimately leading to complete physical dependency and confinement to bed (7).

Mouse models of LD have been pivotal in deciphering disease mechanisms, particularly the role of LB accumulation in neurodegeneration, neurological impairment, and increased seizure susceptibility (11–15). These models emphasise the critical role of energy metabolism in the brain, connecting impaired glycogenolysis to heightened neuronal excitability and reduced seizure thresholds (16). Modified genes in LD mouse models are associated with extensive neurodegenerative changes, including autophagy defects, enlarged lysosomes, β deposits, and mitochondrial anomalies (17). Despite the absence of amyloid plaques in LD patients (18), age-dependent intraneuronal Aβ accumulation has been found in laforin-deficient mice. This finding suggests a critical role of Aβ accumulation in LD pathology, particularly concerning neuronal aberrant excitability (17).

The interaction between epilepsy and neurodegeneration remains a subject of intensive research. Emerging studies suggest that seizures can exacerbate neuronal damage and cognitive decline, highlighting an urgent need for effective treatments to prevent neurodegenerative outcomes. This research revealed a potential role of Aβ accumulation in worsening learning disabilities, primarily linked to neuronal hyperexcitability (19,20). Additionally, a compelling reciprocal relationship exists between Aβ accumulation and abnormal neuronal excitability, creating a vicious cycle that may contribute to neurodegeneration. These dynamics are supported by both preclinical and clinical studies, indicating that mechanisms driving neurodegeneration in Aβ-related conditions may also play a role in epilepsy. Such mechanisms, which escalate neuronal activity, could harm synaptic function and disrupt neural network dynamics (20–22).

However, the early abnormalities that trigger learning disabilities in LD remain to be elucidated. This gap underscores the critical need for further research into the electrophysiological disruptions that may underpin synaptic communication (23). Understanding the early pathological and molecular mechanisms of LD could unveil potential therapeutic targets and establish an optimal intervention window.

According to previous studies, among LD models, the knock-in mouse model *Epm2a^R240X^*, which bears the R240X mutation analogous to the human R241X mutation, displays the most severe epileptic phenotype (15,24). Recent studies on 12-month-old *Epm2a^R240X^* mice have shown increased epileptic-like activity and diminished PTZ-induced seizure thresholds, alongside aberrant synaptic plasticity in the dentate gyrus (DG). Using the *Epm2a^R240X^* model as a precise tool to probe specific aspects of LD, we investigated electrophysiological alterations across different ages in the DG of these mice to unravel the early molecular mechanisms underlying this pathology and that drive its progression. Network hyperexcitability was identified as an early event in the pathophysiology of LD in this model, preceding the deposition of either LBs or amyloid plaques. Amyloid plaque formation in the DG, which may exacerbate the pathophysiology and progression of LD, could be linked to both neuronal aberrant excitability and synaptic anomalies.

In this scenario, treatments like CBD, which can reduce hyperexcitability, hold promise for addressing synaptic plasticity deficits in LD models. CBD is a non-psychoactive compound among *Cannabis sativa* derivatives that has been reported to be well tolerated and safe for clinical use (25,26). Given its anti-seizure properties and favourable clinical profile, CBD is currently approved for the treatment of drug-resistant forms of childhood epilepsy, such as Dravet syndrome and Lennox-Gastaut syndrome, as well as for severe epileptic encephalopathies of childhood onset (26–28). We found that treatment of hippocampal slices with CBD reduced hyperexcitability while rescuing synaptic plasticity in the DG of *Epm2a^R240X^* mice, highlighting the strategic importance of mitigating overexcitability in the management of LD.

## Materials and Methods

### Animals

The *Epm2a^-/-^*, *Epm2b^-/-^* and *Epm2a*^R240X^ mouse models of Lafora disease were generated as previously reported (11,13,15). Mice colonies were bred in the Animal Facility Service of the Instituto de Investigación Sanitaria-Fundación Jiménez Díaz, where they were housed in isolated cages in a 12:12 light/dark cycle at constant temperature (23 °C), with free access to food and water. Procedures were performed in accordance with the Institutional Animal Care and Use Committee-approved protocol of Cincinnati Children’s Hospital and Medical Center. For electrophysiological recordings mice were then transferred to Centro di Ricerca Preclinica of Perugia University and housed in the same conditions described above. All procedures involving animals were performed in conformity with the European Directive 2010/63/EU, and following protocols approved by the Animal Care and Use Committee at the University of Perugia (authorization number 08/2018-UT).

### PAS-diastase staining and immunofluorescence

Mice were transcardially perfused with 4% phosphate-buffered paraformaldehyde. Brains were extracted, sectioned into coronal blocks, dehydrated, and embedded in paraffin. These blocks were subsequently sliced into 5-μm-thick serial sections.

PAS-D staining was conducted using the PAS Kit (Merck, Darmstadt, Germany) and α-amylase from porcine pancreas (Merck, Darmstadt, Germany). Subsequently, sections were counterstained with hematoxylin solution, Gill No. 3 (Merck, Darmstadt, Germany).

For immunofluorescence, sections underwent rehydration in decreasing concentrations of alcohols and antigen retrieval was performed by incubation in 0.1 M sodium citrate buffer (pH 6) at 95 °C. The primary antibodies used were nestin (10 µg/ml, R&D Systems, Minneapolis, Minnesota, USA, Cat #AF2736) and the antibody 4G8 against Aβ (1:250 dilution; Merck, Darmstadt; Germany; Cat. #MABN10). The secondary antibodies were coupled with Alexa Fluor 488 (donkey anti-goat, diluted at 1:400; ThermoFisher Scientific, Waltham, Massachusetts, USA, Cat. #A32814) and Alexa Fluor 594 (donkey anti-mouse, diluted at 1:400; Abcam, Cambridge, UK, Cat. #ab150108).

Samples from 4-6 mice per group were used. To ensure reproducibility, two consecutive sections per mouse were stained and evaluated. Images were captured using a Leica DMLB 2 microscope (Leica, Wetzlar, Germany) connected to a Leica DFC320 FireWire digital microscope camera (Leica, Wetzlar, Germany) and using Zeiss Axioscope 5 (Zeiss, Jena, Germany) connected to an Axiocam 208 color camera (Zeiss, Jena, Germany). LBs, nestin-positive cells and Aβ plaques were quantified using ImageJ by two researchers, with reported values representing the mean of these measurements.

### Drugs

Bicuculline methiodide, picrotoxin and cannabidiol (CBD) were purchased from Tocris Biosciences (Bristol, UK). Drugs were diluted in the appropriate solvent (water for bicuculline, ethanol for picrotoxin and DMSO for CBD) and diluted 1:1000 into the final solution for the desired concentration before conducting the experiments. Drugs were bath applied during the electrophysiological recordings by switching the solution to the one containing the required compounds. The final concentration of ethanol or DMSO (0.1%) did not significantly alter *per se* the electrophysiological parameters investigated.

### Electrophysiology

#### Brain slicing

Mice were sacrificed by cervical dislocation. The brain was collected and immersed in ice-cold artificial cerebrospinal fluid (ACSF) containing (in mM): 126 NaCl, 2.5 KCl, 1.2 MgCl2, 1.2 NaH2PO4, 2.4 CaCl2, 10 glucose, and 25 NaHCO3, continuously bubbled with 95% O2 and 5% CO2, pH 7.4. Transverse hippocampal slices (400 μm or 270 μm thick for extracellular or patch-clamp recordings, respectively) were obtained using a vibratome (LEICA, VT 1200S) with iced ACSF solution for the cutting duration. The slices were then transferred in a recovery chamber with oxygenated ACSF at 30 °C for 30 minutes and then at room temperature (RT) for 1-2 more hours before experimental recordings. Each slice was then transferred into the recording chamber and submerged in ACSF at a constant rate of 2.9–3 mL/min at a temperature of 29 °C.

#### Patch-clamp recordings

Whole-cell patch-clamp recordings (access resistance 10–15 MΩ; holding potential −70 mV) were performed with a Multiclamp 700B amplifier (Molecular Devices, CA) and borosilicate glass pipettes pulled by a P-97 Puller (Sutter Instruments). Recording pipettes were filled with the K^+^-gluconate-based internal solution containing (in mM): 145 K+-gluconate, 0.1 CaCl2, 2 MgCl2, 0.1 EGTA, 10 HEPES, 0.3 Na-GTP and 2 Mg-ATP, adjusted to pH 7.3 with KOH. Pipette resistances ranged from 4 to 7 MΩ. Membrane currents were continuously monitored and access resistance, measured in voltage-clamp mode, was in the range of 10–30 MΩ. Membrane capacitance and resistance of DG granule cells were taken online from the membrane seal test function of pClamp 10.7 (−5 mV step, 10 msec). Injection of depolarizing and hyperpolarizing current steps (1200 msec, 50 pA increments) were used to obtain current-voltage curves and AP numbers at suprathreshold responses. Depolarizing current steps of increasing amplitudes (5 pA increments) were used to determine the AP threshold. When recording spontaneous excitatory postsynaptic currents (sEPSCs), Picrotoxin (50 μM) was added to the ACSF to block GABAA currents. Neurons were clamped at the holding potential (Vh) of −70 mV. Data were acquired with pClamp 10.7 (Molecular Devices), currents were filtered at 0.1 kHz, digitized at 200 μs using Clampex 10.7, analyzed offline using the automatic detection and subsequently checked manually for accuracy.

#### Extracellular recordings

For extracellular recordings, the stimulating electrode was inserted into perforant path fibers and the recording electrode, made of borosilicate glass capillaries filled with 2 M NaCl (resistance 10–15 MΩ), was placed into the in the Dentate Gyrus (DG) close to the granular layer. Stimuli of 0.1Hz, 10 ms duration, and 20–30V amplitude evoked field excitatory post-synaptic potentials (fEPSPs) that in the DG included a population spike (PS) that was 50% of maximum amplitude. The PS amplitude was defined as the average of the amplitude from the peak of the early positivity to the peak negativity and of the amplitude from the peak negativity to the peak late positivity. Axoclamp 2B amplifier (Molecular Devices, USA) was used for extracellular recordings. Traces were filtered at 3 KHz, digitized at 10KHz and stored in a PC. To induce long-term potentiation (LTP) in the DG hippocampal region, a high-frequency stimulation (HFS) protocol, consisting of three trains of 1 s (5min intervals) was delivered at 100 Hz (Kleschevnikov et al 2004) after acquiring a stable baseline for 10 min.

#### Epileptic-like activity and epileptic threshold

Epileptic-like activity in hippocampal DG was induced by perfusing slices with an Mg^2+^-free external solution, to remove the magnesium block from N-methyl-D-aspartate (NMDA) glutamate receptors, in the presence of bicuculline to antagonize gamma-aminobutyric acid A (GABAA) receptors (29,30). The epileptic-like activity was measured both as the mean number of the population spikes and as PS amplitude %, as previously reported (15). A time course of the amplitude of the population spikes in percentage is thus obtained. A similar protocol, removing bicuculline from the solution, can be used to assess epileptic threshold.

### Biochemistry

#### Western Blotting (WB) experiments

Tissues were homogenized at 4◦C in an ice-cold buffer containing 0.32 M sucrose, 0.1 mM phenylmethylulfonyl fluoride (PMSF), 1 mM HEPES, 1 mM MgCl, and 1 mM NaF, supplemented with protease inhibitors (Complete™, Sigma-Aldrich, St. Louis, MI, USA) and phosphatase inhibitors (PhosSTOP™, Sigma-Aldrich). Protein samples were separated using a denaturing sodium dodecyl-sulfate polyacrylamide gel electrophoresis (SDS-PAGE) followed by transfer onto nitrocellulose membranes. The membranes were incubated for 1 h at RT in blocking solution (I-block, Tris-buffered saline [TBS] 1X, 20% Tween 20) on a shaker. The membranes were then incubated with the specific primary antibodies in blocking solution overnight at 4◦C, and the following day, after three washes with TBS and 0.1% Tween 20 (TBSt), they were incubated with the corresponding horseradish peroxidase (HRP)-conjugated secondary antibody in blocking solution for 1 h at RT. After washing with TBSt, the membranes were developed using electrochemiluminescence (ECL) reagents (Bio-Rad, Hercules, CA, USA). Finally, membranes were scanned using a Chemidoc (Bio-Rad) with Image Lab software (Bio-Rad). The protein bands were quantified using computer-assisted imaging (Image Lab, Bio-Rad). Protein levels were expressed as relative optical density (OD) measurements, normalized to tubulin, and expressed as a percentage of the control mean. The following primary antibodies were used: rabbit anti-GluA1 (WB 1:1000, #13185S, Cell Signaling, Danvers, MA, USA), mouse anti-GluA2 (WB 1:2000, #75-002, Neuromab, UC Davis, Davis CA), mouse anti-GluA3 (WB 1:1000, #MAB5416, Millipore, Millipore, Burlington, MA, USA), monoclonal rabbit anti-phosphoSer845-GluA1 (WB 1:1000, #ab76321,Abcam), rabbit anti-GluN2A (WB 1:1000, #M264, Sigma-Aldrich), rabbit anti-GluN2B (WB 1:1000, #14544s, Cell signaling, Danvers, MA, USA), rabbit anti-ERK (WB 1:1000, #9102, Cell Signaling, Danvers, MA, USA), rabbit anti-phosphoERK (WB, 1:1000, #9101, Cell Signaling, Danvers, MA, USA) mouse anti-tubulin (WB 1:10000, #T9026, Sigma-Aldrich). The following secondary antibodies were used for WB analysis: goat anti-rabbit HRP and goat anti-mouse HRP (#1706515 and #1706516, respectively, Bio-Rad).

### Statistical analysis

Data analysis for electrophysiological recordings and histological assays were performed offline using Clampfit 10.7 (Molecular Devices) and GraphPad Prism 9.0 (GraphPad Software, Inc., La Jolla, CA 92037, USA) software. Values given in the text and figures are expressed as mean ± S.E.M. The “n” denotes the number of cells or field potential recorded for electrophysiological analysis, as well as the number of animals used for histological approaches. Two-way ANOVA, Student’s t-test, or the Mann-Whitney test were used for statistical analysis. The significance level was set at *p<0.05. Since no differences were found in the basal membrane properties of DG granule cells, epileptic-like activity and synaptic plasticity between young (1-3 months) and old (6-12 months) WT mice (data not shown), the data from these groups of animals were pooled together for analysis.

## Results

### Electrophysiological characterisation of *Epm2a^-/-^*, *Epm2b^-/-^,* and *Epm2a^R240X^* mouse models at 12 months in the DG

Electrophysiological characterisation was performed in the DG of 12-month-old *Epm2a^-/-^*, *Epm2b^-/-^,* and *Epm2a^R240X^*mice to assess which genotype presents the most severe electrophysiological phenotype, which could then be used as a model for more in-depth, time-dependent investigations. We previously reported that *Epm2a^R240X^* knock-in mice exhibit earlier cognitive decline and enhanced susceptibility to PTZ-induced seizures compared to *Epm2a^-/-^* mice (15). Therefore, epileptic-like activity, epileptic threshold, and LTP were analysed in the DG of these three LD models. Epileptic-like activity in hippocampal slices was induced in both LD and age-matched WT mice by applying an inducing protocol (see Methods section) able to elicit only slight epileptic-like activity in WT mice. Our data show that epileptic-like activity is similar in *Epm2a^-/-^* and *Epm2a^R240X^* mice, and significantly higher than in wild-type (WT) mice (Fig. S1A). Under the same experimental conditions, *Epm2b^-/-^* mice displayed milder epileptic-like activity, which, however, was significantly different from both WT and laforin-deficient mice (Fig. S1A). Evaluation of the epileptic threshold in this brain region revealed a comparable threshold in *Epm2b^-/-^*and WT mice, while both laforin-deficient genotypes displayed lower epileptic thresholds compared to WT (Fig. S1B). Nevertheless, the time course graph shows a significant increase in the PS amplitude in *Epm2a^R240X^*compared to *Epm2a^-/-^*, occurring within the first 10 minutes of 0Mg^2+^ solution superperfusion (Fig. S1B). This finding indicates that the DG of *Epm2a^R240X^* mice have a lower epileptic threshold than that of *Epm2a^-/-^* mice.

LTP in the DG of the three genotypes was then investigated, revealing that the LTP of DG granule cells was impaired in *Epm2a^-/-^* mice, preserved at physiological levels in *Epm2b^-/-^* mice, and significantly enhanced in *Epm2a^R240X^* mice compared to WT mice, as previously described (15).

Taken together, these results indicate that laforin-deficient genotypes display an electrophysiological phenotype of similar severity, albeit with opposite deficits in synaptic plasticity, which will require further investigation. *Epm2a^R240X^*, however, shows a lower epileptic threshold than the *Epm2a^-/-^* genotype. This is consistent with data obtained *in vivo* following treatment of the animals with PTZ (15). It can therefore be concluded that the more severe histopathological and behavioural phenotype observed in the *Epm2a^R240X^* model (15) is underpinned by an electrophysiological phenotype characterised by enhanced network hyperexcitability.

### Age-dependent alterations of neuronal basal membrane properties in the DG of *Epm2a*^R240X^ **mice**

To investigate possible changes in the intrinsic electrical membrane properties in DG granule cells of *Epm2a^R240X^* mice, patch-clamp recordings were performed in hippocampal slices from 1-, 3-, and 12-month-old *Epm2a^R240X^* mice and age-matched WT animals. DG cells from *Epm2a*^R240X^ animals presented a significant increase in membrane resistance, a significant reduction in membrane capacitance, and a reduced time constant as early as 3 months (Fig. 1A). Furthermore, analysis of the current-voltage relationship and resting membrane potentials (V_rest_) also revealed significant age-dependent differences between the groups. Specifically, reduced inward currents, elicited at hyperpolarising voltage steps, and a hyperpolarised V_rest_, were observed in *Epm2a*^R240X^ animals as early as 3 months of age compared to WT (Fig. 1B).

**Figure 1.**
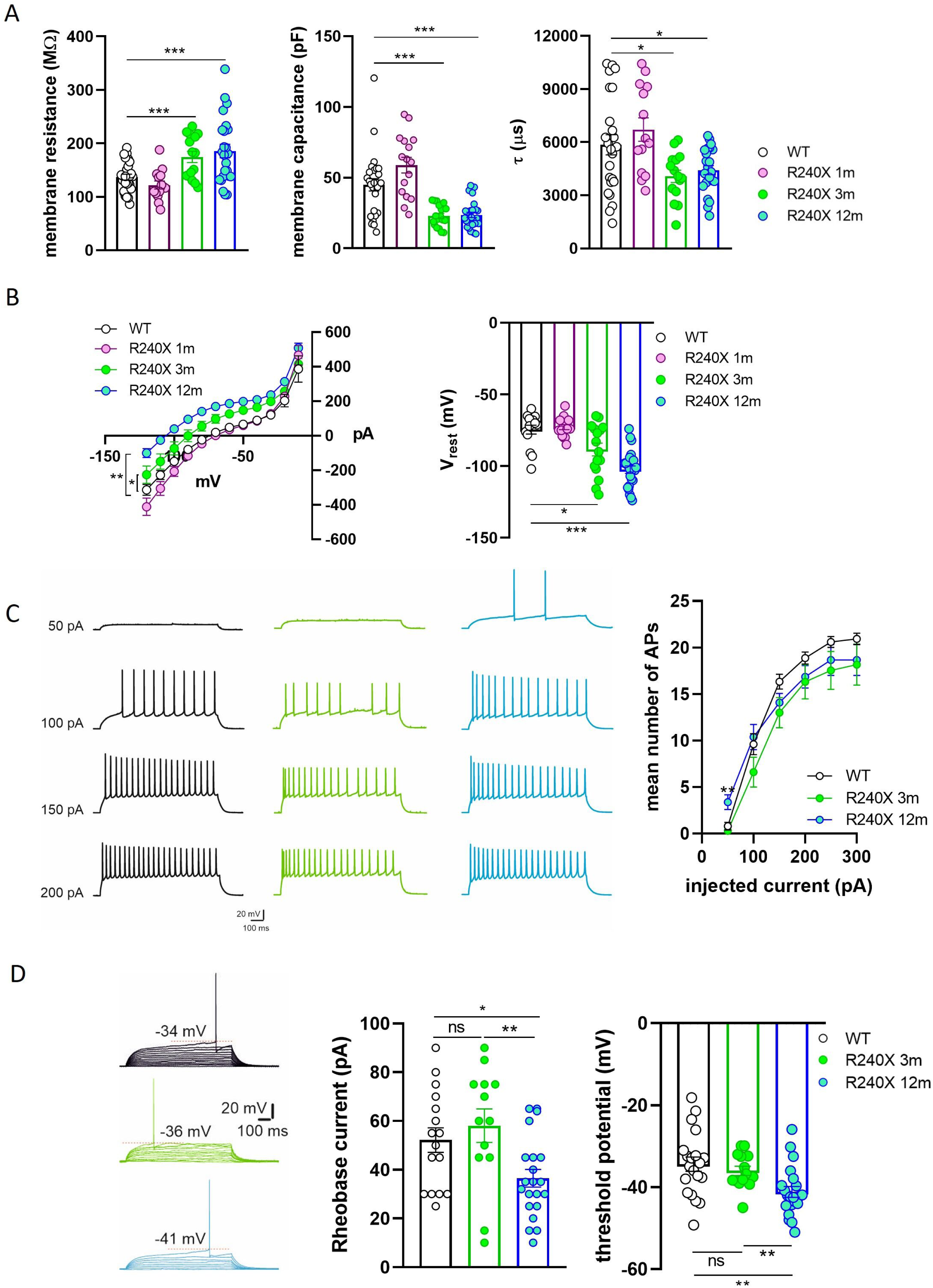
Intrinsic membrane properties of DG granule cells in 1-, 3- and 12-month-old *Epm2a*^R240X^ mice. **A)** Histograms showing significative age-dependent increase of membrane resistance (Rm) and reduction of membrane capacitance (Cm) and time constant (τ) (**Rm:** WT=136.9 ± 5.37 MΩ; 1m-R240X= 122.2 ± 7.54 MΩ, n>10; 3m-R240X= 174.6 ± 10.5 MΩ; 12m-R240X= 185.6 ± 12.7 MΩ; **Cm:** WT = 44.9 ± 4.33 pF; 1m-R240X= 58.8 ± 5.53 pF; 3m-R240X = 22.9 ± 1.85 pF; 12m-R240X = 23.67 ± 1.83 pF; **τ:** WT: 5847±571 μs; 1m-R240X: 6708 ± 656 μs; 3m-R240X: 4067±346 μs; 12m-R240X: 4413±254 μs; n>10, *p<0.05, ***p<0.001 Student’s t-test). **B)** Mean current–voltage plot showing significant age-dependent alterations in *Epm2a*^R240X^ mice (*p<0.05 WT vs 3m-R240X; **p<0.01 3m-R240X vs 12m-R240X; ***p<0.001 WT vs 12m-R240X; two-way ANOVA). Right: histogram highlighting the age-dependent hyperpolarization of the resting membrane potential in *Epm2a^R240X^* mice (WT: -74.56 ± 3.04 mV; 1m-R240X: -72.87 ± 1.75 mV; 3m-R240X: -88.44 ± 4.45 mV; 12m-R240X: -102.5 ± 2.61 mV; n>10, *p<0.05; **p<0.01; ***p<0.001; Student’s t-test). **C)** Patterns of action potential (AP) firing in WT (black) and *Epm2a^R240X^* mice at 3 (green) and 12 (cyan) months. The plot representing the mean number of AP shows a significant difference in the firing pattern discharge elicited by the first step of depolarizing current injected between WT and 12-month-old *Epm2a*^R240X^ mice (WT: 0.61 ± 0.4, n>10; 3m-R240X: 0.3 ± 0.1, n>10; 12m-R240X: 1.9 ± 0.8, n=10; **p < 0.01, WT vs 12m-R240X, Student’s t-test). **D)** Current–clamp recordings (5 pA–stepped depolarizing current injections; 1 sec), scaled to show AP threshold (red dotted line), in WT (black) and *Epm2a*^R240X^ mice at 3 (green) and 12 (cyan) months. Histograms showing a significant reduction of the rheobase current (middle, WT: 52.19 ± 5.04, n>10; 3m-R240X: 58.08 ± 6.8; 12m-R240X: 36.48 ± 3.6; n>10, *p<0.05 WT vs 12m-R240X; **p<0.01, 3m-R240X vs 12m-R240X Student’s t-test) and a hyperpolarized threshold potential (right, WT: -34.4 ± 1.76; 3m-R240X: -36.02 ± 1.14; 12m-R240X: -41.16 ± 1.33; n>10, **p<0.01, Student’s t-test) in 12-month-old *Epm2a*^R240X^. Values are reported as mean ± SEM.

Membrane excitability was also evaluated by applying depolarising and hyperpolarising current steps. Action potential (AP) numbers were similar in *Epm2a^R240X^* and WT mice, except for the first depolarising current step that triggered significantly higher APs in 12-month-old *Epm2a^R240X^* animals (Fig. 1C). In line with this, the rheobase current was reduced, while the threshold potential was hyperpolarised, only in 12-month-old *Epm2a^R240X^* animals (Fig. 1D). Notably, all intrinsic electrical membrane properties of DG granule cells are preserved at physiological levels in 1-month-old *Epm2a^R240X^* mice.

Analysis of the spontaneous excitatory postsynaptic currents (sEPSCs) in DG granule cells revealed no differences between WT and *Epm2a^R240X^*animals (Fig. S2).

### Epileptic-like activity and aberrant LTP in 3-month-old *Epm2a*^R240X^ mice

Since 12-month-old *Epm2a^R240X^*mice have been reported to display aberrant excitability of DG granule cells and epileptic-like activity (15), we considered exploring earlier time points in this model to identify more precocious events underlying the progression of LD.

Epileptic-like activity was induced in hippocampal slices of 1-, 3-, and 12-month-old *Epm2a^R240X^* mice and age-matched controls. The PS number and amplitude were significantly increased in all ages of *Epm2a^R240X^* mice compared to WT animals, in an age dependent manner, indicating more intense epileptic-like activity in 12-month-old *Epm2a^R240X^* mice relative to younger mice (Fig. 2A). Interestingly, epileptic-like activity was comparable between 1- and 3-month-old *Epm2a*^R240X^ mice, suggesting that network hyperexcitability is a very early event in the progression of this pathology, potentially preceding the deposition of LBs (Fig. S3), at least in this model.

**Figure 2.**
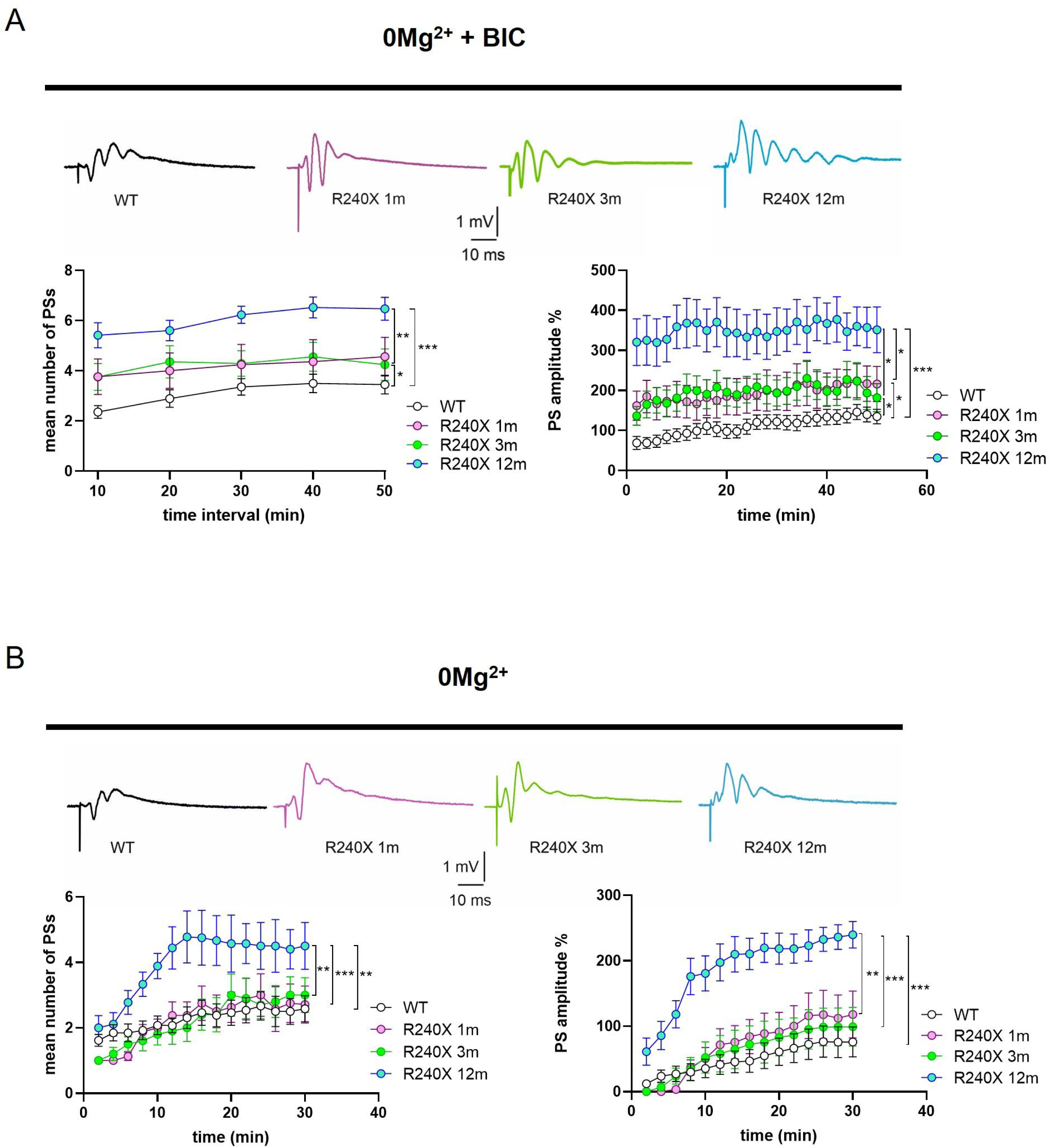
Epileptic–like activity in DG slices of 1-, 3- and 12-month-old *Epm2a*^R240X^ mice. **A)** Representative traces of FPs (upper panel) and time-course graph (lower panels) of the mean number (left) and amplitude (right) of PSs recorded in the DG of WT (black), 1- (pink), 3- (green) and 12-month-old (cyan) *Epm2a^R240X^*mice in a magnesium-free ACSF in the presence of 0.1 µM bicuculline, showing a time-dependent increase of the epileptic-like activity in LD mice (**PS number:** WT: 3.4 ± 0.4, n>10; 1m-R240X: 4.5 ± 0.8, n=5; 3m-R240X: 4.2 ± 0.6, n=10; 12m-R240X: 6.5 ± 0.4, n>10; **PS amplitude:** WT: 135 ± 17.9 %, n>10; 1m-R240X: 216 ± 34.8 %, n=10; 3m-R240X: 182 ± 34.8 %, n=10; 12m-R240X: 351 ± 57.0 %, n>10). **B)** Representative traces of FPs (upper panel) and time-course graph (lower panels) of the mean number (left) and amplitude (right) of PS measured in the DG of WT (black) and *Epm2a*^R240X^ mice at 1 (pink), 3 (green) and 12 (cyan) months of age, in a magnesium-free ACSF in the absence of bicuculline, showing the lower epileptic threshold in 12-month-old *Epm2a^R240X^* mice (**PS number:** WT: 2.6 ± 0.4, n>10; 1m-R240X: 2.7 ± 0.6, n=7; 3m-R240X: 3.0 ± 0.53, n=8; 12m-R240X: 4.5 ± 0.72, n=6; **PS amplitude:** WT: 75.8 ± 22.9 %, n=10; 1m-R240X: 118 ± 35.8 %, n=8; 3m-R240X: 99.1 ± 29.5 %, n=10; 12m-R240X: 240 ± 20.3 %, n=11). Data are reported as means ± SEM. *p<0.05, **p<0.01, ***p<0.001, two-way ANOVA.

To better dissect age-dependent differences, epileptic threshold parameters were analysed in 1- and 3-month-old *Epm2a^R240X^* mice (see Methods section), measuring PS number and amplitude. We previously reported a lower epileptic threshold in 12-month-old *Epm2a*^R240X^ mice compared to WT (15). No significant differences were found between young *Epm2a^R240X^* and WT mice (Fig. 2B), indicating that the lower epileptic threshold is a characteristic displayed only by older *Epm2a^R240X^* mice. Taken together, these data suggest that the DG of *Epm2a^R240X^* mice may already exhibit altered network excitability as early as 1 month, although to a lesser extent than in their older counterparts.

Since Aβ peptides have been shown to be deeply involved in neuronal hyperexcitability associated with synaptic and network dysfunction in models of various neurodegenerative disorders (20), we hypothesised that amyloid plaque formation in the DG might exert a role in the pathophysiology of LD. A significant increase in 4G8-positive deposits was found in the DG of 12-month-old *Epm2a^R240X^* compared to WT mice (Figs. 3A and B).

**Figure 3.**
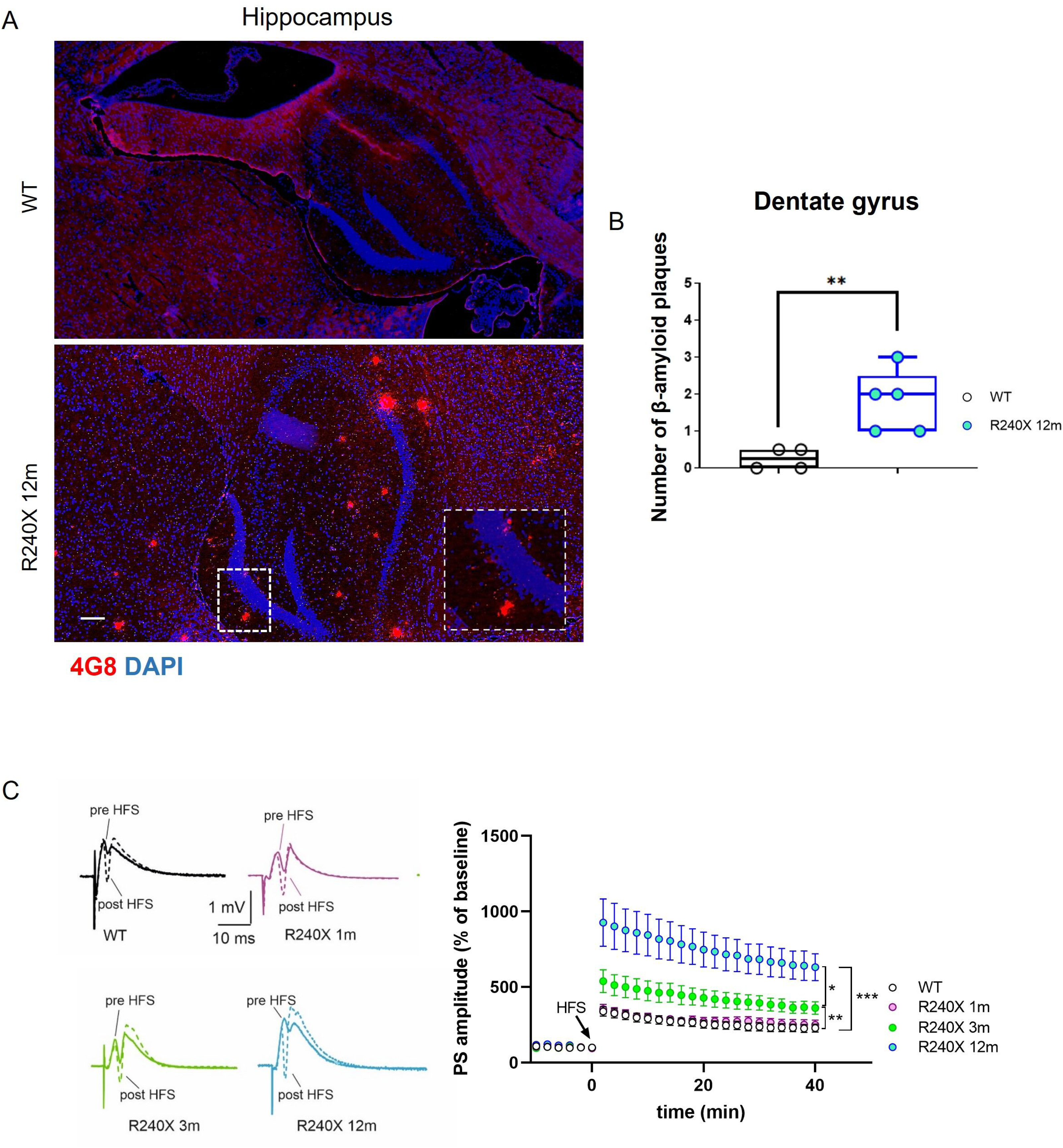
Quantification of amyloid-β plaques and time-dependent LTP alteration in the DG of WT and *Epm2a^R240X^* mice. **A)** Immunofluorescence using the 4G8 antibody revealed abundant presence of amyloid-β plaques in the hippocampal region of WT and *Epm2a^R240X^* mice. **B)** Quantitative comparison of amyloid deposits in the DG at 12 months of age. Data are shown as median of independent samples. Whiskers in box plots indicate the minimum and maximum values. For statistical analyisis, a non-parametric Mann Whitney test was carried out. Scale Bar = 100µm. n=4-5 mice per genotype. **C)** Representative PSs traces recorded before (continuous line) and 40 min after (dotted line) the HFS protocol in DG slices of WT (black), 1- (pink), 3- (green) and 12-month-old (cyan) *Epm2a^R240X^* mice. The time–course plot of PSs amplitudes recorded in the DG before and after HFS protocol shows a time-dependent hyper-plasticity in LD mice. Note that 1-month-old *Epm2a^R240X^* mice display a physiological LTP (WT: 230 ± 26.4 %, n>10; 1m-R240X: 250 ± 31.4 %, n=5; 3m-R240X: 437 ± 52.4 %, n>10; 12m-R240X: 632 ± 89.3 %, n=9). Data are reported as means ± SEM. *p<0.05, **p<0.01, ***p<0.001, two-way ANOVA.

The LTP of DG granule cells was therefore investigated in this brain region. We previously reported abnormally increased LTP in the DG of 12-month-old *Epm2a^R240X^* mice compared to WT mice (15). This aberrant LTP is likely due to neuronal hyperexcitability (hLTP) and is associated with cognitive and learning impairments (15). *Epm2a^R240X^*mice at 3 months also displayed an increase in LTP compared to WT (Fig. 3C). The comparison between 3- and 12-month-old knock-in mice showed a significant reduction in LTP amplitude at 3 months. Interestingly, LTP in the DG of 1-month-old *Epm2a^R240X^* mice is preserved at physiological levels. These results, together, suggest that hyperexcitability precedes synaptic deficits as well as the deposition of either LBs and Aβ, at least in this model of LD.

Given the role of ionotropic receptors in synaptic plasticity, the expression levels of AMPA and NMDA receptor subunits were analysed in total homogenates prepared from hippocampal DG areas. As shown in supplementary Figs. 4A and 4B, WB analysis revealed no alteration in the protein levels of the GluN2A and GluN2B subunits of NMDA receptors or the GluA1, GluA2, and GluA3 subunits of AMPA receptors in *Epm2a^R240X^* compared to WT mice. In addition, evaluation of the phosphorylation levels (pThr845) of the GluA1 subunit, known to be relevant for receptor anchoring at the postsynaptic membrane (31), and phosphorylation of the signalling protein ERK (pERK/ERK) showed no significant change (Fig. S4C).

**Figure 4.**
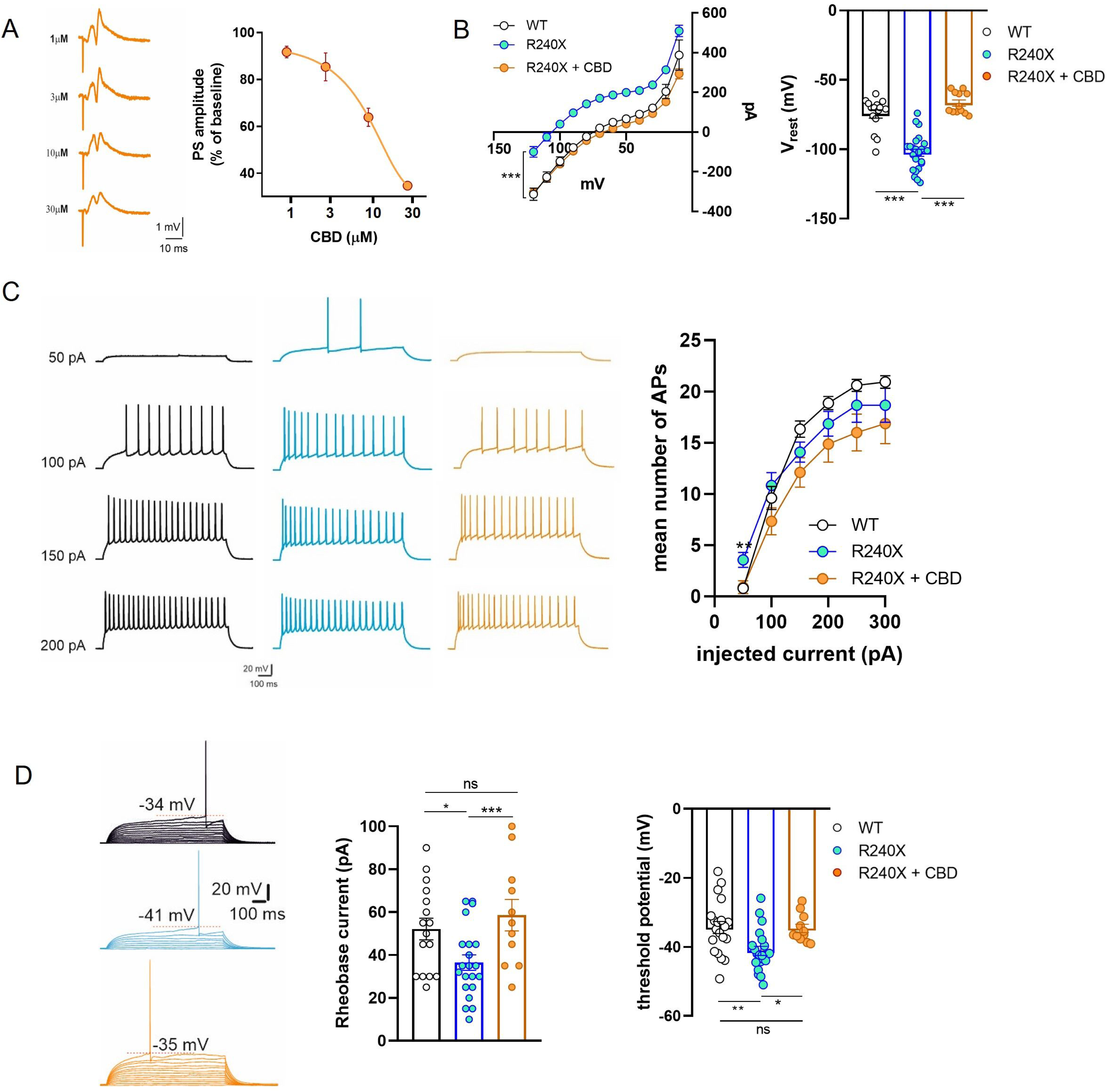
Effect of CBD on 12-month-old *Epm2a^R240X^* mice DG granule cells excitability. **A)** Effect of CBD on physiological synaptic transmission in the DG region. Left: representative traces of PSs recorded on increasing doses of CBD. Dose-response curve showing that CBD 3 μM is not able to alter the physiological synaptic transmission (CBD 1 μM: 91.7±2.5 % of baseline; CBD 3 μM: 85.4±5.9 % of baseline; CBD 10 μM: 63.8± 3.8 % of baseline; CBD 30 μM: 34.6±0.7 % of baseline; IC50: 10.96 ± 0.08 nM). **B)** CBD 3 μM restores the physiological current-voltage relationship in the granule cells of 12-month-old *Epm2a^R240X^* mice (middle, ***p<0.001, R240X vs R240X + CBD, two-way ANOVA) and normalizes the hyperpolarized resting membrane potential (right, WT= -74.5 ± 3 mV; R240X= -102.5 ± 2.6 mV; R240X+CBD= -66.7 ± 2.2 mV; n>10, ***p<0.001, Student’s t-test). **C)** Patterns of AP firing of DG granule cells in WT (black) and *Epm2a^R240X^* mice in the absence (cyan) and in the presence of 3μM CBD (orange). Note that CBD is able to recover the firing pattern discharge elicited by the first step of depolarizing current in 12-month-old *Epm2a^R240X^* mice (WT: 0.6 ± 1.4, n>10; R240X: 3.6 ± 0.7, n>10; R240X+CBD: 0.9 ± 0.6, n=10; **p<0.01, Student’s t-test). **D)** Representative current–clamp recordings (5 pA–stepped depolarizing current injections; 1 sec), scaled to show AP threshold (red dotted line), in DG granule cells from WT (black) and *Epm2a^R240X^*mice in the absence (cyan) and in the presence of 3μM CBD (orange). Histograms showing that 3µM CBD is able to normalize the rheobase current (WT: 52.2 ± 5 pA; 12m-R240X: 36.5 ± 3.6 pA; R240X+CBD: 58.6 ± 7.3 pA; n>10, *p<0.05, ***p<0.001 Student’s t-test) and the threshold potential (WT: -34.4 ± 1.7 mV; R240X: -41.2 ± 1.3 mV; R240X+CBD: -34.7 ± 1.2 mV; n>10, *p<0.05, **p<0.01, Student’s t-test) to physiological levels in 12-month-old *Epm2a*^R240X^ mice. Data are reported as means ± SEM.

We then investigated whether the lower epileptic threshold and the larger amplitude of LTP elicited by perforant path synaptic inputs could result from increased hippocampal neurogenesis (32,33). To this end, nestin-expressing progenitor cells (34) were labelled in the hippocampus of WT and *Epm2a^R240X^* mice at 3 and 12 months of age (Fig. S6A). Quantification revealed that *Epm2a^R240X^* mice exhibit an increased number of progenitor cells in all hippocampal regions compared to WT mice at 12 months, but not at 3 months (Fig. S5B-G).

### Reduction of neuronal hyperexcitability rescues LTP in the DG of 12-month-old *Epm2a*^R240X^ **mice**

We then assessed whether reducing hyperexcitability with CBD could effectively counteract aberrant synaptic plasticity in this LD model at 12 months. First, a dose-response curve was established to evaluate the effects of CBD on synaptic transmission, aiming to determine the optimal CBD concentration. After acquiring a stable PS response for 10 min, CBD was bath applied for 20 min at concentrations ranging from 1 to 30 μM. As expected, CBD dose-dependently reduced the PS amplitude response compared to baseline, with maximal effect at 30 μM (Fig. 4A). The highest CBD dose that did not affect physiological transmission (3 μM) was then tested for its ability to rescue the electrophysiological alterations and the epileptic-like activity observed in the DG region of 12-month-old *Epm2a^R240X^* mice. As shown in Fig. 4B, CBD restored the current-voltage relationship and the resting membrane potential of *Epm2a^R240X^* mice granule cells to control levels. CBD also reduced the number of action potentials in *Epm2a*^R240X^ animals (Fig. 4C) by normalising the rheobase current and the threshold potential (Fig. 4D). Moreover, CBD completely restored the epileptic-like activity to control levels, both in the presence and absence of bicuculline (Figs. 5A and 5B). When investigating the effects of CBD on synaptic plasticity, treatment with CBD was seen to rescue the aberrant LTP observed in the DG of *Epm2a^R240X^* mice to control values (Fig. 5C). Taken together, these data demonstrate that enhanced excitability of DG granule cells underlies aberrant synaptic plasticity in this region and that counteracting neuronal hyperexcitability may be a valuable strategy to rescue LTP.

**Figure 5.**
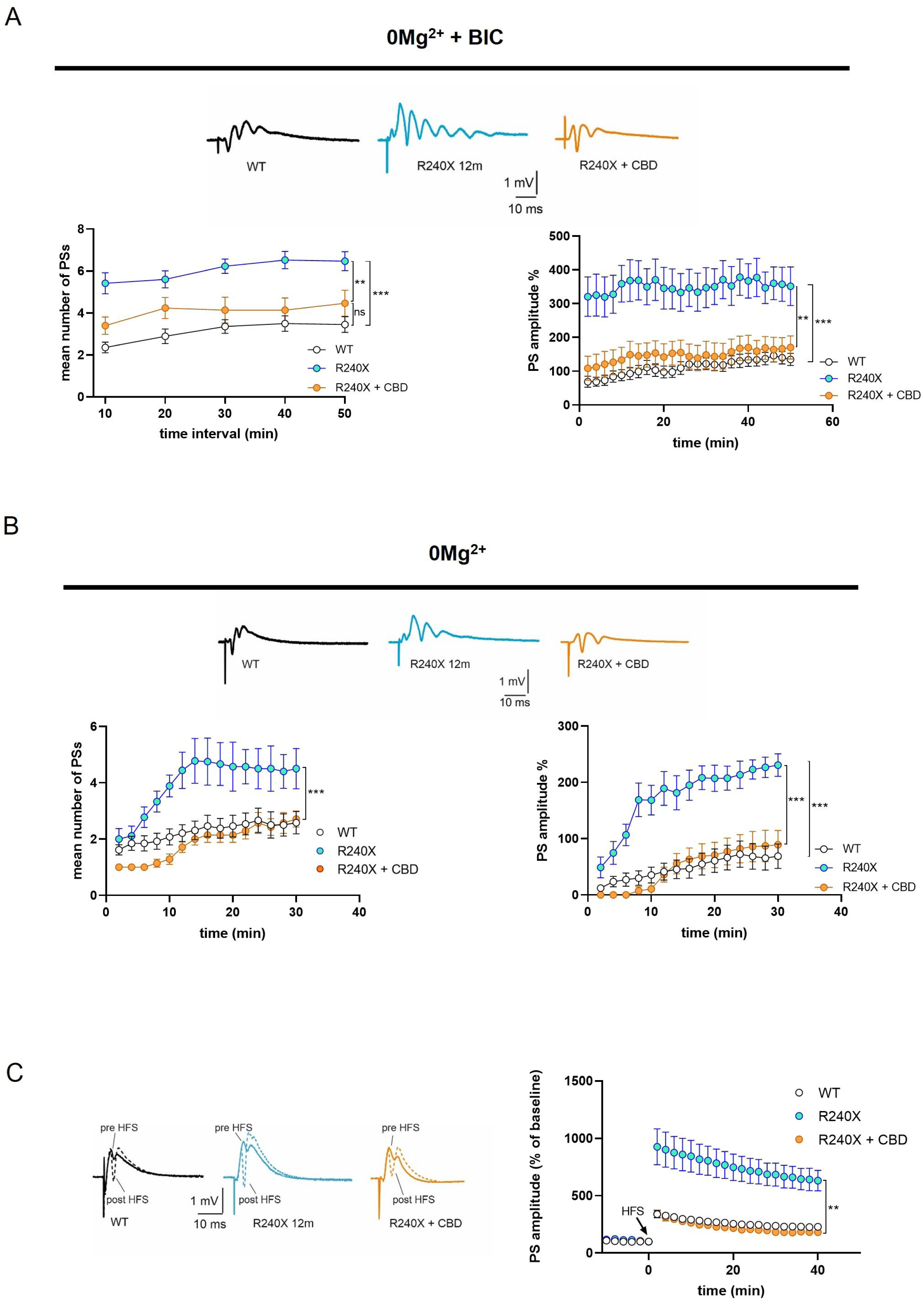
Effect of CBD on epileptic-like activity and LTP in the DG of 12-month-old *Epm2a^R240X^* mice. **A)** Representative traces and time-course graph of FPs recorded in magnesium-free + 0.1 μM bicuculline ACSF solution in the DG of WT (black traces) and 12-month-old *Epm2a*^R240X^ mice in the absence (cyan traces) or in the presence (orange traces) of 3µM CBD. Note that 3µM CBD is able to recover the mean number and amplitude of PSs in 12-month-old *Epm2a*^R240X^ mice to control levels (**PS number:** WT: 3.4 ± 0.4, n>10; R240X: 6.5 ± 0.4, n>10; R240X+CBD: 4.5 ± 0.6, n=6; **PS amplitude:** WT: 135 ± 17.9 %, n>10; R240X: 351 ± 57.0 %, n>10; R240X+CBD: 170 ± 34.3 %, n=7; **p<0.01, ***p<0.001, two-way ANOVA). **B)** Representative traces and time-course graph of FPs recorded in magnesium-free ACSF solution in the DG of WT (black traces) and 12-month-oldd *Epm2a^R240X^* mice in the absence (cyan traces) or in the presence (orange traces) of 3µM CBD. Note that 3µM CBD is able to recover the mean number and amplitude of PSs, in 12-month-old *Epm2a^R240X^* mice to control levels (**PS number:** WT: 2.6 ± 0.4, n>10; R240X: 4.5 ± 0.72, n=6; R240X+CBD: 2.7 ± 0.28, n=7; **PS amplitude:** WT: 75.8 ± 22.9 %, n=10; R240X: 240 ± 20.3 %, n=11; R240X+CBD: 89.2 ± 25.3 %, n=7; ***p<0.001, two-way ANOVA). **C)** Representative traces and time-course graph of PSs recorded before (continuous line) and 40 min after (dotted line) the HFS protocol in WT (black), 12-month-old *Epm2a^R240X^* (cyan) and in *Epm2a^R240X^* mice in the presence of 3μM CBD (orange). Note that treatment with 3μM CBD is able to restore the aberrant LTP observed in the DG of *Epm2a^R240X^* to physiological levels (WT: 230 ± 26.4 %, n>10; R240X: 632 ± 89.3 %; R240X + CBD: 182 ± 15.4 %; **p<0.01, two-way Anova). Data are reported as means ± SEM.

## Discussion

This study outlines key findings demonstrating that network hyperexcitability is an early event in the pathophysiology of LD, manifesting prior to the deposition of LBs and Aβ, and preceding the onset of synaptic deterioration (Fig. 6). Furthermore, our research highlights progressive changes in synaptic plasticity within the DG of the *Epm2a^R240X^* model, marked by early alterations in neuronal excitability, a pattern observed in various other neurodegenerative disorders. These findings suggest that intervention with targeted treatments during the early stages of the disease, potentially before the onset of neurological clinical symptoms in humans, could significantly influence disease progression. Therefore, identifying and capitalising on this early therapeutic window is crucial for effectively modifying the course of the disease.

**Figure 6.**
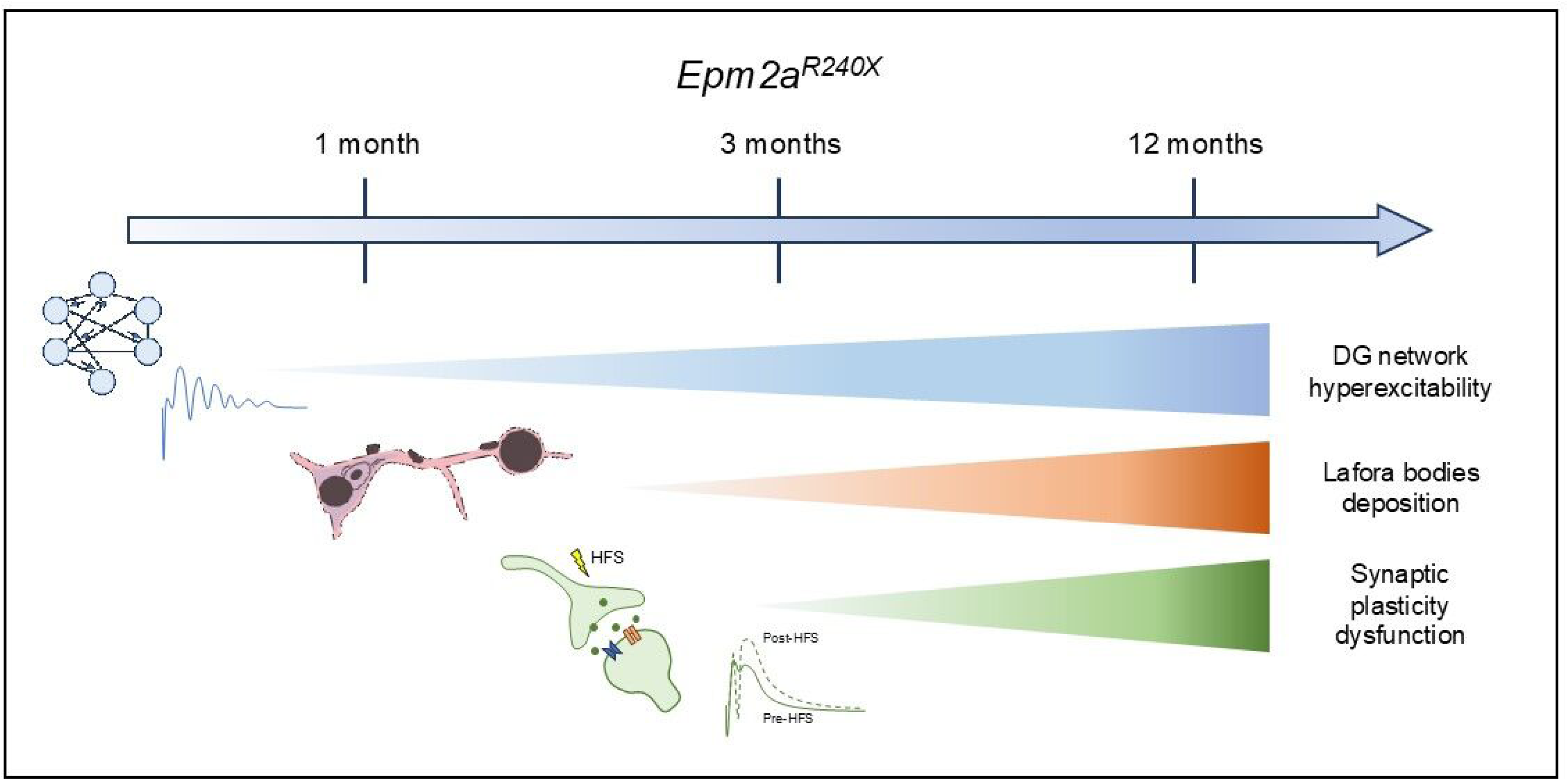
Network hyperexcitability as an early event in LD progression. Our results suggest that network hyperexcitability might represent one of the early pathogenic steps in LD, preceding synaptic deterioration and clear LBs or Aβ deposition. The schematic representation of LBs was made taking inspiration from the original drawing of LBs made by Lafora (8).

The mechanisms underlying aberrant excitability and synaptic dysfunction in the onset and progression of LD remain unknown. Indeed, neurodegeneration and seizure susceptibility in LD have been correlated with LB deposition (6,14,35). However, more recent evidence demonstrates abnormalities in dendritic spines and related cognitive-behavioural deficits occurring in LD mouse models long before the appearance of LBs (23), suggesting possible alterations in synaptic communication that could constitute an early substrate for the neuronal altered excitability underlying the epileptogenic event and cognitive impairment. The complex bidirectional relationship between epilepsy and cognitive decline remains a matter of scientific debate. Clinical and preclinical evidence suggests that epilepsy and cognitive decline are closely intertwined (19,22) and may share common underlying pathophysiological mechanisms (20,21) that still need to be fully elucidated. In this scenario, it is possible to hypothesise that hyperexcitability is the *primum movens* of cognitive decline in LD as well.

In support of this hypothesis, our previous work demonstrated enhanced excitability and lower epileptic threshold, along with aberrant hyperplasticity (hLTP) in the DG of 12-month-old *Epm2a^R240X^* mice, while the CA1 region remained unaffected (15). This is in agreement with previous studies reporting that increased excitability drives enhanced synaptic transmission and LTP magnitude in the DG (36–38), and that this synaptic dysfunction spreads spatiotemporally, starting from the DG and extending to the CA1 (37). Thus, hyperexcitability and neurodegeneration appear to be a continuum encompassing epilepsy and cognitive decline, with aberrant excitatory neuronal activity triggering compensatory mechanisms that lead to a loss of homeostatic plasticity, in turn contributing to network dysfunction in a self-powering loop (19,20). In this scenario, the aberrant form of LTP that we observed in the DG of older *Epm2a^R240X^*mice seems to be based on hyperexcitability and could be defined as hLTP (15).

To gain a deeper understanding of the mechanisms underlying the early stages of the disease, we investigated whether the alterations observed in the DG of 12-month-old *Epm2a^R240X^*mice were already present in younger (1- and 3-month-old) mice. We first examined potential age-dependent changes in the intrinsic membrane electrical properties underlying heightened excitability in DG granule cells of *Epm2a^R240X^*mice. Reduced rheobase current and hyperpolarised threshold potential were only found in 12-month-old *Epm2a*^R240X^ mice, while in younger *Epm2a*^R240X^ mice these parameters were comparable to those in WT mice. In line with this, we also observed an increased number of APs elicited by the first step of depolarising current injection in 12-month-old *Epm2a^R240X^* mice. Although this evidence indicates that enhanced excitability in DG granule cells develops only in the advanced stages of the disease, other parameters were altered as early as 3 months. Specifically, analysis of the current-voltage relationship and resting membrane potential revealed age-dependent changes in *Epm2a^R240X^* mice, characterised by a progressive reduction in inward current amplitudes recorded at hyperpolarised membrane potentials, along with a hyperpolarised V_rest_. The hyperpolarised resting potential likely reflects differential control by multiple voltage-gated channels, such as sodium, calcium, potassium, and HCN channels (39–42). In particular, *h* currents are deeply involved in modulating excitatory postsynaptic potential (EPSP) duration, thereby contributing to temporal summation and the regulation of neuronal excitability (43,44). We cannot rule out the possibility that at least some of these conductances are altered in the *Epm2a^R240X^* model, as observed in other types of epilepsy (45–49), but further investigation will be needed to clarify this issue. Moreover, evidence of the hyperpolarised resting potential, coupled with a reduced rheobase current in older *Epm2a^R240X^*mice, suggests the possibility of a unique complex system whose behaviour results from either altered voltage-dependence and kinetics or an aberrant density or distribution of these channels.

Finally, intrinsic membrane properties, such as membrane capacitance, resistance, and the time constant tau, were already altered in *Epm2a^R240X^*mice at 3 months to the same extent as in the older animals. The reduction in the time constant tau can be considered a precocious indicator of excitability (50,51), since it determines how quickly the membrane potential changes in response to ionic currents. This suggests that DG granule cells in 3-month-old *Epm2a^R240X^* mice are more prone to excitability, and this feature is already evident at this young age.

We previously reported that the DG granule cells of 12-month-old *Epm2a^R240X^*mice display enhanced epileptic-like activity, a lower epileptic threshold, and aberrant synaptic plasticity compared to WT mice (15). Similar forms of enhanced synaptic plasticity have been described in several genetic neuropathologies (52,53) and have been associated with cognitive impairment rather than improvement. The *Epm2a^R240X^* model exhibits cognitive deficits (15) and hyperplasticity, raising the possibility that the aberrant LTP observed in the DG is a result of the hyperexcitability of granule cells. In this study, we investigated the epileptic-like activity and synaptic plasticity in the DG of young *Epm2a^R240X^* mice to assess whether the alterations observed are also present in the early stages of LD. It was found that, although the epileptic threshold is preserved, epileptic-like activity is already increased in both 1- and 3-month-old *Epm2a^R240X^* mice, albeit to a significantly lesser extent than in older animals, demonstrating a time-dependent progression of this parameter in LD. Accordingly, analysis of LTP in the DG of 3-month-old mice showed a significant increase compared to WT, but also a significant decrease compared to 12-month-old *Epm2a^R240X^*mice. On the other hand, the DG of 1-month-old mice displayed physiological LTP. This latter finding demonstrates that network overexcitability is an early event in the progression of LD, preceding both LBs and Aβ deposition, as well as synaptic damage, at least in this model of LD, thereby defining a potential time window for therapeutic intervention.

Interestingly, although time-dependent Aβ deposition has been previously reported in laforin KO mice (17), we demonstrated for the first time the presence of Aβ deposits in the hippocampus of *Epm2a^R240X^* mice. It is plausible that Aβ deposition contributes to the pathophysiology of LD. Indeed, clinical and preclinical studies highlight that Aβ deposition and aberrant neuronal and network excitability are closely intertwined in a vicious cycle, leading to dysfunctional network activity and rearrangements that ultimately result in neurodegeneration (19,20). Another possibility is that altered excitability arises from aberrant neurogenesis in this LD model. We demonstrate an increased number of progenitor cells in all hippocampal regions of *Epm2a^R240X^* compared to WT mice at 12 months of age. A complex interplay between hyperexcitability and aberrant neurogenesis has been described, with seizures increasing neurogenesis in the hippocampus, further contributing to an overactive state due to the abnormal integration of new neurons. In this scenario, newly formed neurons from aberrant neurogenesis may form synaptic connections that enhance network excitability, potentially leading to a vicious cycle of hyperexcitability and further abnormal neurogenesis (54,55).

To further investigate our hypothesis that hLTP could be the consequence of enhanced excitability, excitability was reduced by treating hippocampal slices of 12-month-old *Epm2a^R240X^* mice with CBD, an emerging anti-seizure drug. CBD has recently been proposed as a viable therapeutic substitute for people with seizure disorders that are refractory, such as Lennox-Gastaut syndrome and Dravet syndrome (DS) (26). Compared with traditional anti-seizure drugs, CBD is an effective anticonvulsant with a higher specificity and fewer neurotoxic effects (56). Furthermore, CBD exhibits high selectivity, does not induce excitability in the central nervous system, and can reduce both the duration and amplitude of the post-discharge cAMP response element-binding protein (CREB). (57). CBD has been tested in several epilepsy animal models, including MES (maximal electroshock seizure model), pentylenetetrazol, and pilocarpine (58–61), consistently showing anti-seizure effects. As expected, treatment of DG slices with CBD at a concentration that does not alter physiological transmission *per se* was able to reduce epileptic-like activity and to rescue the epileptic threshold to control levels in 12-month-old *Epm2a^R240X^* mice. The mechanisms through which CBD modulates the excitability of the DG in *Epm2a^R240X^*mice still need to be elucidated but may rely on the multi-target action of CBD. Indeed, more than 50 molecular targets have been identified for CBD (62,63), several of which are ionotropic and metabotropic receptors deeply involved in neuronal function and transmission, as well as in synaptic calcium mobilisation and membrane potential.

Intriguingly, treatment with CBD also rescued LTP in the DG of *Epm2a^R240X^* mice to control levels. This is the first demonstration that reducing excitability can constitute a strategy to rescue synaptic plasticity in a mouse model of LD. Our findings are in line with previous reports showing that the anti-seizure effects of CBD are paralleled by improvements in cognitive and behavioural deficits in the *Scn1a^+/−^*genetic mouse model of DS (61,64).

Taken together, our results shed light on neuronal alterations occurring in the early stages of LD and provide proof of principle that modulation of neuronal excitability can constitute a tool to rescue synaptic dysfunction in this model of LD.

## Aknowledgements

We thank Prof. Paolo Calabresi for critical revising the manuscript, Chiara Galizia for her technical support in biochemical experiments, Francisco Wandosell for his generous gift of 4G8 and nestin antibodies and Rosario Moratalla (Instituto Cajal, CSIC, Madrid Spain) for the kind gift of the TH antibody. We also thank the Animal Facility of Instituto de Investigación Sanitaria-Fundación Jiménez Díaz for their technical assistance, and the “DEFEAT-LD study group” of PNRR-MR1-2022-12376430 - Project ’Drug discovEry and repurposing to Find a trEAtmenT for Lafora Disease (DEFEAT-LD)’.

## Conflict of interest statement

Cinzia Costa has received research funding, speaker honoraria, and travel support from Bial, Eisai, Europe Limited, GW Pharma, Jazz Pharmaceuticals, Lusopharma, PIAM Pharma, and UCB Pharma.

None of these companies had any role in the study design, data collection, analysis or interpretation, manuscript preparation, or the decision to submit the article for publication.

---

**Figure S1.**
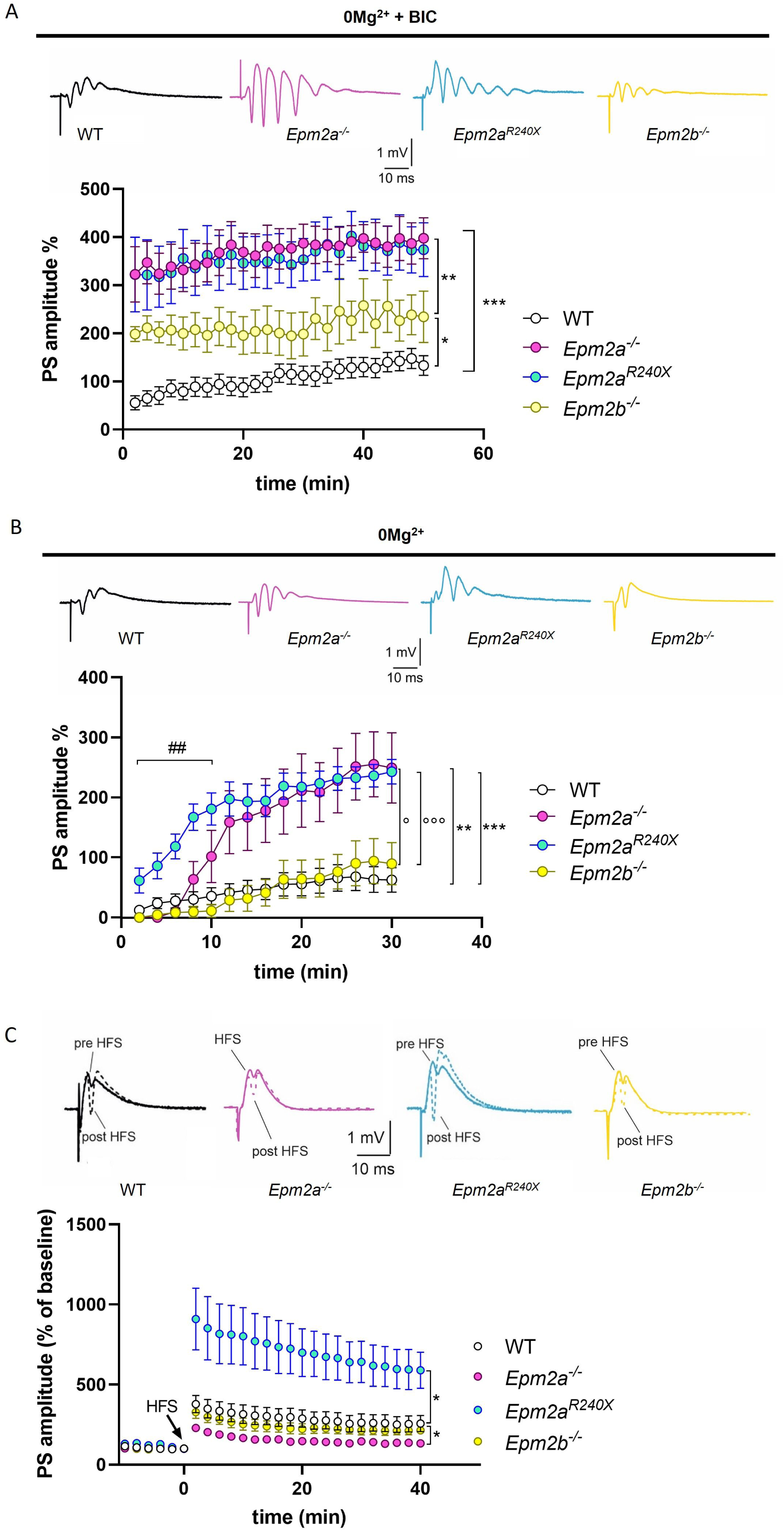
Epileptic-like activity, epileptic threshold and LTP in the DG of 12-month-old *Epm2a*^-/-^, *Epm2b^-/-^* and *Epm2a*^R240X^ mice. **A)** Representative traces of FPs (upper panel) and time-course graph (lower panel) of the mean PS amplitude % recorded in the DG of 12-month-old WT (black), *Epm2a^-/-^* (purple), *Epm2b^-/-^* (yellow) and *Epm2a^R240X^* (cyan) mice in a magnesium-free ACSF in the presence of 0.1 µM bicuculline, showing significative differences between groups (WT: 133 ± 20.7 %, n>10; *Epm2a^-/-^*: 398 ± 42.5 %, n=9; *Epm2b^-/-^*: 235 ± 53.4 %, n=5; *Epm2a^R240X^*: 305 ± 33.1 %, n>10; *p<0.05 WT vs *Epm2b^-/-^*; **p<0.01 *Epm2b^-/-^* vs *Epm2a^-/-^* and *Epm2a^R240X^*; ***p<0.001 WT vs *Epm2a^-/-^* and *Epm2a^R240X^*, two-way Anova). Data are reported as means ± SEM. **B)** Representative traces of FPs (upper panel) and time-course graph (lower panel) of the mean PS amplitude % measured in the DG of 12-month-old WT (black) *Epm2a^-/-^* (purple), *Epm2b^-/-^* (yellow) and *Epm2a^R240X^* (cyan) mice, in a magnesium-free ACSF in the absence of bicuculline, showing a lower epileptic threshold in *Epm2a^-/-^* and *Epm2a^R240X^* mice. (WT: 62.3 ± 20.4 %, n>10; *Epm2a^-/-^*: 249 ± 58.6 %, n=9; *Epm2b^-/-^*: 89.5 ± 35.3 %, n=7; *Epm2a^R240X^*: 243 ± 20.4 %, n>10; °p<0.05 *Epm2b^-/-^* vs *Epm2a^-/-^*; **p<0.01 WT vs *Epm2a^-/-^*; ##p<0.01 *Epm2a^R240X^* vs *Epm2a^-/-^*; °°°p<0.001 *Epm2b^-/-^* vs *Epm2a^R240X^*; ***p<0.001 WT vs *Epm2a^R240X^*; two-way ANOVA). Data are reported as means ± SEM. **C)** Representative PSs traces recorded before (continuous line) and 40 min after (dotted line) the HFS protocol in DG slices of 12-month-old WT (black), *Epm2a^-/-^* (purple), *Epm2b^-/-^*(yellow) and *Epm2a^R240X^* (cyan) mice. The time–course plot of PSs amplitudes recorded in the DG before and after HFS protocol shows significative differences between groups. Note that *Epm2b^-/-^* mice display a physiological LTP (WT: 257 ± 50 %, n=9; *Epm2a^-/-^*: 133 ± 11.1 %, n=6; *Epm2b^-/-^*: 214 ± 23.5 %, n=6; *Epm2a^R240X^*: 590 ± 112.6 %, n=7). Data are reported as means ± SEM. *p<0.05, two-way ANOVA.

**Figure S2.**
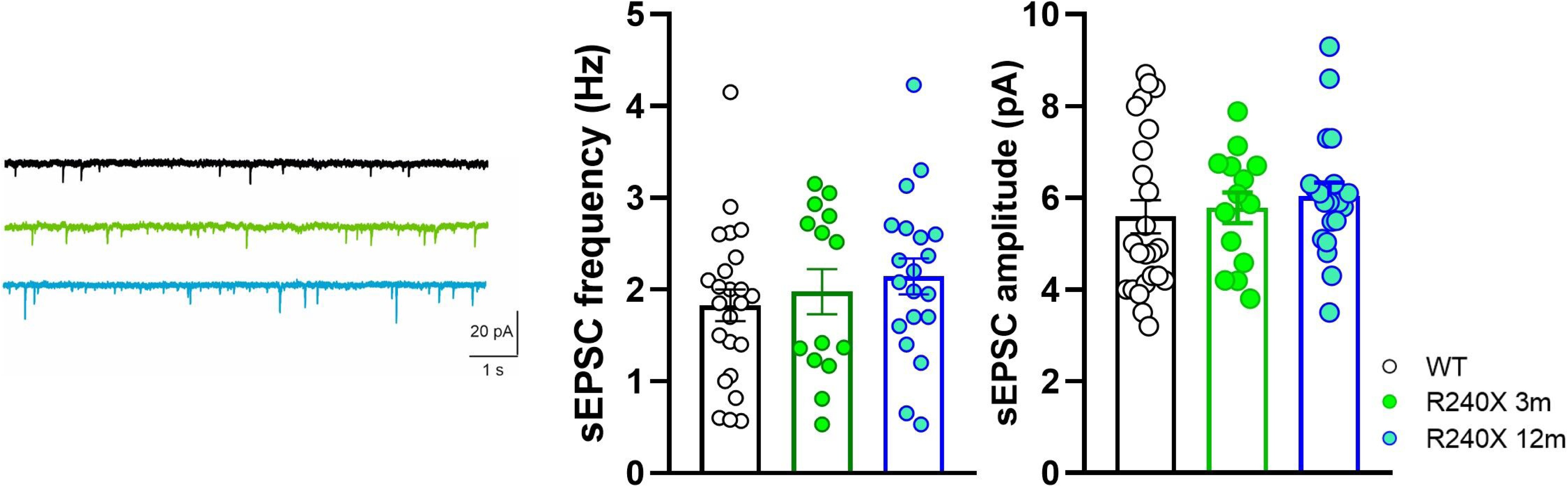
Spontaneous glutamatergic transmission in DG slices of 3- and 12-month-old *Epm2a*^R240X^ mice. **A)** Representative sEPSC traces recorded from DG granule cells of WT (black traces), 3-month-old Epm2a^R240X^ (green traces) and 12-mont-old Epm2a^R240X^ (cyan traces) mice. **B)** Histograms reporting the mean frequency (WT: 1.8 ± 0.17 Hz, n>10; 3m-R240X: 1.9 ± 0.24 Hz, n>10; 12m-R240X: 2.1 ± 0.19 Hz, n>10; ; p>0.05, Student’s t-test) and amplitude (WT: 5.6 ± 0.36 pA, n>10; 3m-R240X: 5.8 ± 0.33 pA, n>10; 12m-R240X: 6 ± 0.3 pA, n>10; p>0.05, Student’s t-test) of the sEPSC show no differences between the groups.

**Figure S3.**
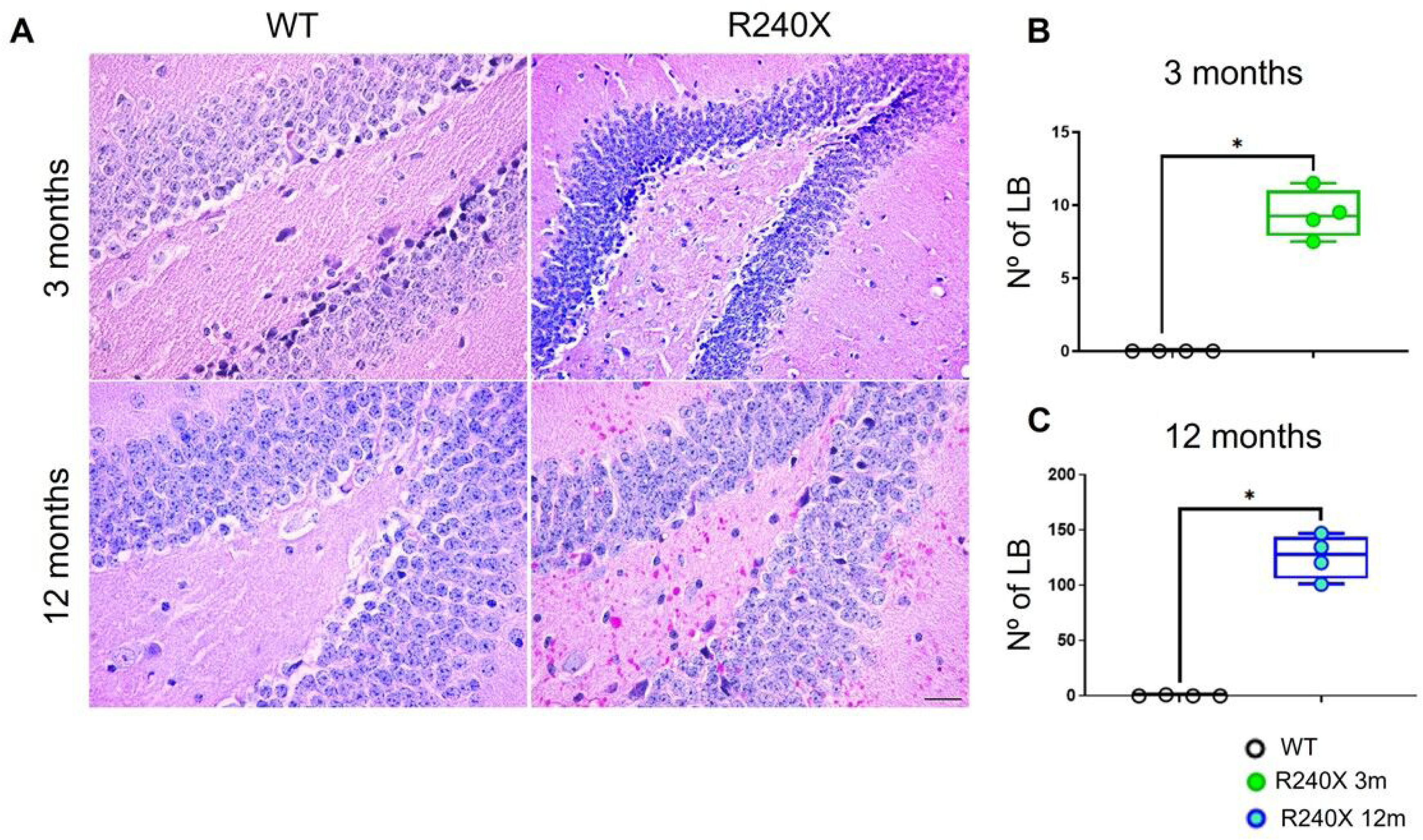
LB formation in the dentate gyrus of WT and *Epm2a^R240X^* mice. **A)** PAS-D staining of the DG of 3 and 12-month-old WT and *Epm2a^R240X^* mice. **B)** Quantitative comparison of LB number in the DG of the hippocampus at 3 (left), and 12 (right) months of age. Data are shown as median of independent samples. Whiskers in box plots indicate the minimum and maximum values. Statistical analysis was performed using a non-parametric Mann Whitney test. * p < 0.05. Scale Bar = 25µm.

**Figure S4.**
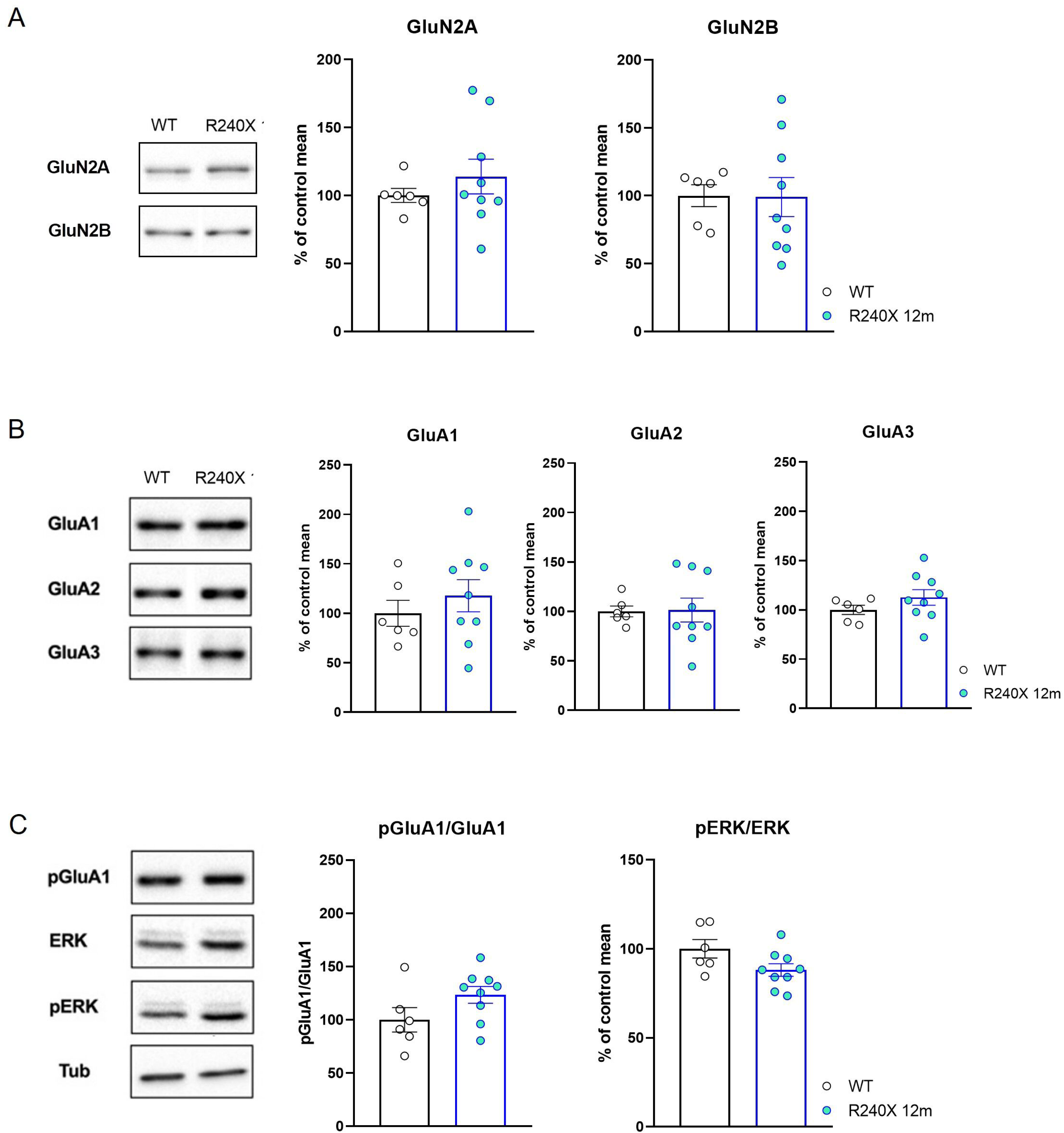
Ionotropic glutamate receptor subunit expression and evaluation of p-GluA1 and pERK in the DG of 12-month-old *Epm2a^R240X^* and WT mice. **A-B)** Western blotting representative images (left) and bar graphs (right) of densitometric quantification of GluN2A and GluN2B subunits of NMDA receptors (A) and GluA1, GluA2 and GluA3 subunits of AMPA receptors (B) in total cell homogenates prepared from DG of 12-month-old WT and *Epm2a^R240X^* mice. **C)** Western blot representative images (left) and bar graphs (right) of densitometric quantification of phospho-Thr845-GluA1 (pGluA1) and phospho-ERK/ERK (pERK/ERK) in total cell homogenates prepared from DG of WT and *Epm2a^R240X^* mice. Tubulin was used for normalization. Data are expressed as percent of the mean value of protein levels in WT group (N=6,9).

**Figure S5.**
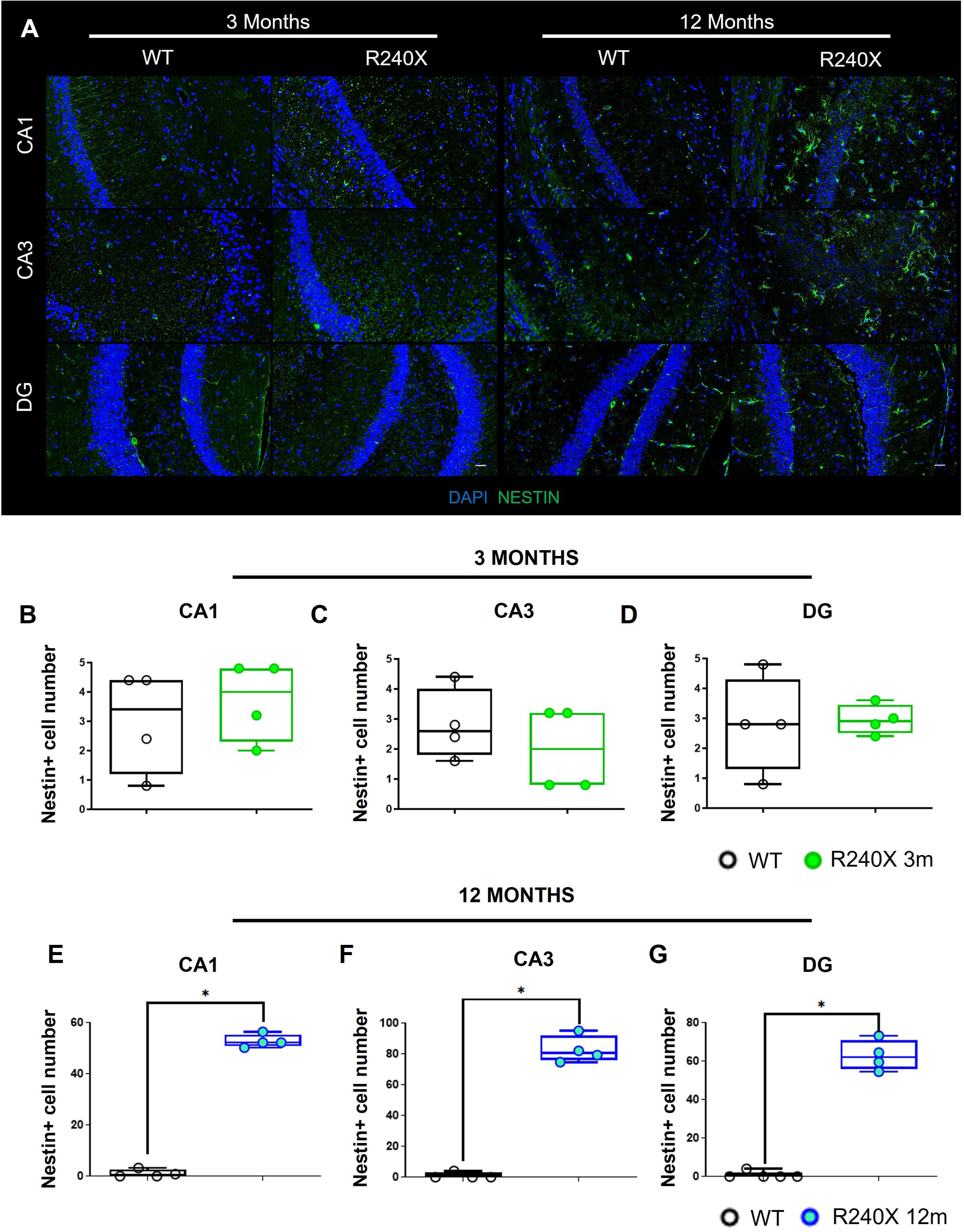
Nestin-expressing progenitor cells in the hippocampus of WT and *Epm2a^R240X^* mice. **A)** Representative immunofluorescence images of nestin in hippocampal regions CA1, CA3 and DG at 3 and 12 months of age. **B-G)** Quantification of nestin-positive cells in the CA1 (**B and E**), CA3 (**C and F**) and DG **(D and G)** hippocampal regions at 3 (upper panel) and 12 (lower panel) months of age. Data are shown as median of independent samples. Whiskers in box plots indicate the minimum and maximum values. Statistical analysis was conducted using a non-parametric Mann Whitney test. * p < 0.05; ** p < 0.001. Scale Bar = 25µm. n= 4 mice per genotype.

